# Association of lanthipeptide genes with TnpA_REP_ transposases in marine picocyanobacteria

**DOI:** 10.1101/2020.03.09.984088

**Authors:** Raphaël Laurenceau, Nicolas Raho, Zev Cariani, Christina Bliem, Sean M. Kearney, Elaina Thomas, Mohammed A. M. Osman, Sallie W. Chisholm

## Abstract

Lanthipeptides are a family of ribosomally synthesized, post-translationally modified peptides that are widespread among bacteria, typically functioning as antibacterials. The marine picocyanobacteria *Prochlorococcus* and *Synechococcus* produce an unusual and diverse set of lanthipeptides of unknown function called prochlorosins. While well-studied model bacteria produce one or two different molecules of this type, a single picocyanobacterium can produce as many as 80; the community of picocyanobacteria in a single milliliter of seawater can collectively encode up to 10,000 prochlorosins. The molecular events that led to this expansion and diversification of the lanthipeptide repertoire in picocyanobacteria – the numerically dominant photosynthesizers in the oceans – is unknown.

We present evidence for an unusual association between prochlorosin genes with a single-stranded DNA transposase belonging to the TnpA_REP_ family. The genes co-occur and co-localize across the phylogeny of marine picocyanobacteria forming a distinct association pattern within genomes, most likely resulting from the transposase activity. Given the role of TnpA_REP_ homologs in other bacteria, we propose - based on genomic structures - that they contribute to the creation of the prochlorosin structural diversity through a diversifying recombination mechanism.

**Post-submission note:** Since the original submission of this manuscript to bioRxiv, we have refined our phylogenetic analysis of the TnpA-REP transposases and we do not find strong evidence for or against a causal relationship between the presence of TnpA-REP transposases and the expansion and diversification of lanthipeptides genes. While the genetic association described in the manuscript remains valid, adjacent TnpA-REP and Prochlorosins do not appear phylogenetically linked, which might simply be the consequence of high rates of recombination for both genes. More genomic data are needed to untangle the driving force behind the lanthipeptide gene expansion in marine picocyanobacteria.

**IMPORTANCE:** Only a few mechanisms have been described that promote the diversification of a targeted gene region in bacteria. We present indirect evidence that the TnpA_REP_ transposases associated with prochlorosins in picocyanobacteria could represent a novel such mechanism, and explain the extreme expansion and diversification of prochlorosins in this abundant marine microbe.

## OBSERVATION

Cubillos-Ruiz et al. [1] revealed that prochlorosins – lanthipeptides produced by marine picocyanobacteria – represent an extreme expansion and diversification of the lanthipeptide family described in other bacteria [2]. While typical model bacteria encode 1 or 2 of these compounds, select strains of picocyanobacteria encode up to 80 different ones – representing an extensive diversity of cyclic peptide structures. The unusual abundance of these genes in the otherwise very streamlined genomes of these tiny cells suggests a strong selection pressure to maintain and diversify them [1]. Their biological function in the ecosystem is unknown.

Prochlorosin biosynthesis involves the production of precursor peptides (ProcA) which are matured by a promiscuous modifying enzyme (ProcM). ProcA peptides are composed of an N-terminal signal peptide – the “leader peptide” – directing the sequence to ProcM which cyclizes the C-terminal “core peptide” by adding intramolecular lanthionine bonds, the defining feature of lanthipeptides [2]. The leader peptide is subsequently cleaved, yielding the final cyclic prochlorosin. While the leader peptide-encoding sequence is well-conserved across picocyanobacteria, the core peptide-encoding sequence is responsible for the immense diversity of prochlorosins, through the accumulation of small insertion/deletion events [1]. The precise molecular events leading to this unique expansion and diversification has remained elusive.

A likely mechanism for the expansion of prochlorosins might be diversifying recombination – a process by which a localized region of DNA (with respect to a given gene) is modified through a dedicated recombination mechanism, leading to a hotspot of sequence diversity across closely related bacterial cells. Diversification can lead to variability in gene expression, or structural changes in a particular protein domain – and the selective advantage it provides is often (but not exclusively) related to ‘camouflage’, like avoiding infection by phages or avoiding detection by the immune system. Precise mechanisms of diversifying recombination that have been described in other bacteria are: 1) localized homologous recombination (bacteriocins, like colicins [3], [4] or syringacins [5], capsule serotypes [6], adhesins [7], or restriction-modification systems [8]; 2) invertase-mediated shuffling (pilins [9], capsule serotypes and various surface-exposed antigens [10], [11]; 3) diversity-generating retroelements (transcriptional regulators, pilin-like proteins and signaling proteins [12]; and 4) integrons (shuffling of genes cassettes including antibiotic resistance and other virulence genes [13], [14]. We conducted BLAST searches of a vast prochlorosin database for the machinery operating in these four other bacterial systems and found nothing similar.

In the process of our detailed exploration of prochlorosin genomic structure and context, however, we observed the association of prochlorosin genes with a single-stranded DNA transposase from the TnpA_REP_ family [15]–[18] (Fig. 1). TnpA_REP_ transposases are single-stranded DNA recombinases that – unlike the closely-related IS200-like transposases – are considered domesticated in the sense that they do not move across genomes in a typical transposon structure, but rather are involved in creating large repeat regions containing ‘REPs’ (Repetitive Extragenic Palindromes, motifs that form DNA hairpin structures) [15], [18]–[20]. For example, there are 600 REP repeats in *E. coli*, representing ∼1% of its genome [21].

**FIG 1.**
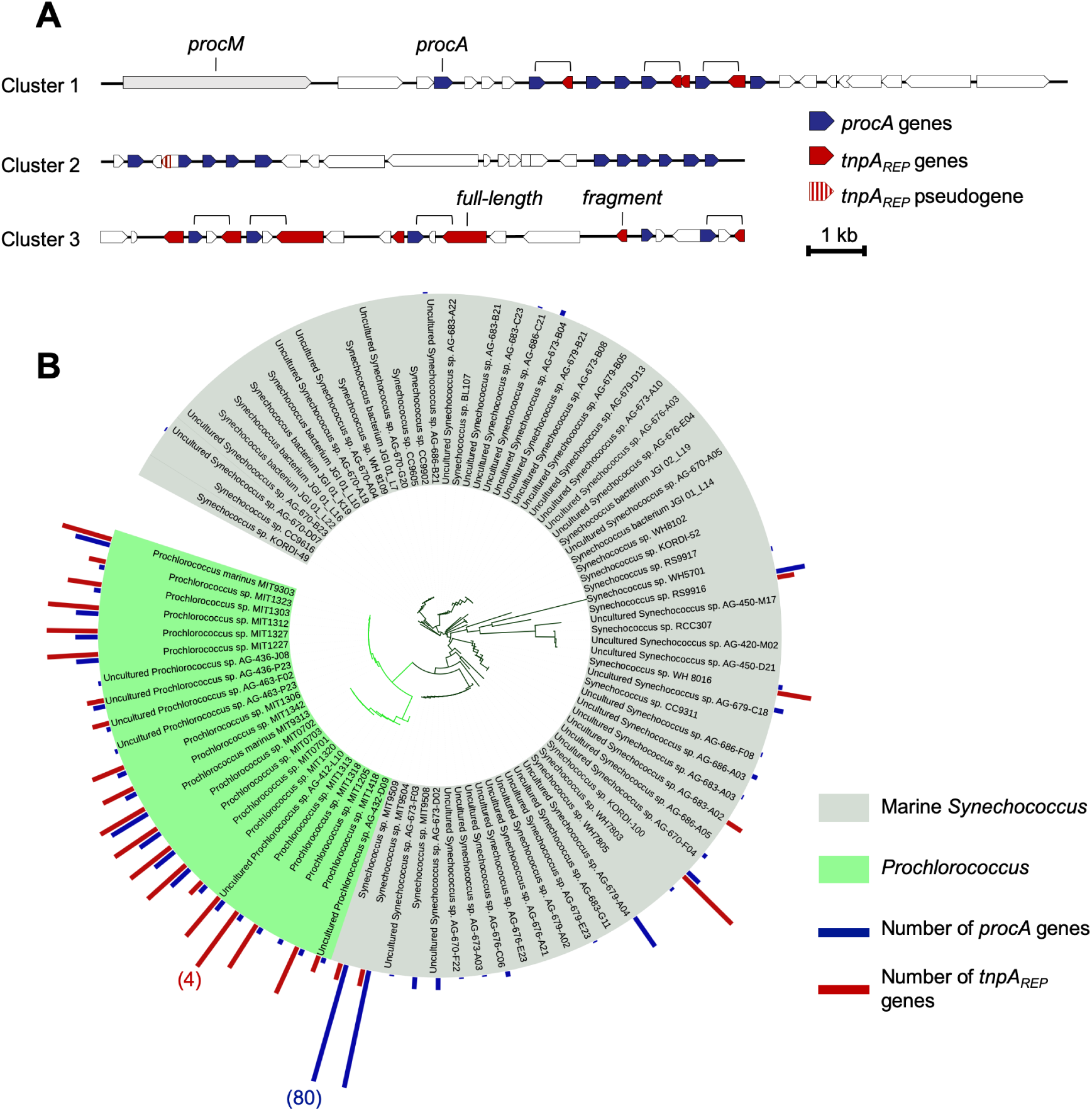
**(A)** Example of co-localization of *tnpA*_*REP*_ and *procA* genes in several prochlorosin clusters in the *Prochlorococcus* MIT9313 genome. Prochlorosin genes in blue were annotated following [1]. Brackets indicate the repeated arrangement pattern of *procA* genes facing *tnpA*_*REP*_ genes. **(B)** Co-occurrence of *procA* and *tnpA*_*REP*_ genes in marine picocyanobacteria. The phylogenetic distance tree was built using the built-in IMG-Proportal phylogeny tool and the tree was annotated using the Interactive Tree Of Life (iTOL). *Prochlorococcus* genomes are restricted to the LLIV clade, as all other clades are lacking both *tnpA*_*REP*_ genes (not shown) and Prochlorosins [1]. The bar diagram on the tree indicates the number of individual *procA* genes (blue) and individual *tnpA*_*REP*_ genes (only ‘full-length’ genes encoding for > 200 amino acids TnpA_REP_, in red) within a given strain (no bar indicates the absence of the genes). Bracketed numbers on the red and blue bars indicate scale.

Importantly, *tnpA*_*REP*_ genes are physically associated with REP repeat regions (usually flanking them), and several studies have confirmed a biochemical link between TnpA_REP_ and REP motifs, supporting their role as the primary driver for the expansion and diversification of REP regions in bacterial genomes [16], [17], [22], [23], a feature that is reminiscent with the pattern of sustained expansion, diversification, and elimination of *procA* genes we have observed in marine picocyanobacteria [1].

We found a number of TnpA_REP_ family transposase homologs in marine picocyanobacteria, all of them forming a monophyletic group that appears to be descended from TnpA_REP_ in other cyanobacteria such as *Trichodesmium erythraeum* (Sup Fig 1, 2). They belong to a subclass of TnpA_REP_ (RAYT group 1 in [18], or subclass 2.1 in [24]) which possess an additional conserved domain in the C-terminal region, likely involved in DNA-binding [24]. In marine picocyanobacteria, the *tnpA*_*REP*_ genes tend to co-localize with clusters or individual *procA* genes, which encode for the peptides that are cyclized by ProcM to become the final product: prochlorosins (Fig. 1A). Out of the 136 *tnpA*_*REP*_ sequences (including full-length copies and fragments) found in marine picocyanobacteria, 32 % are within 200 bp of a *procA* gene (likely an underestimate because repeat-containing regions of the genome are difficult to assemble and contigs often end next to transposases genes, Sup Fig 3). Full-length transposase copies – as well as numerous copies of truncated fragments (often corresponding to the C-terminal extremity of TnpA_REP_, Sup Fig 4) – are interspersed with *procA* gene clusters (Fig. 1A, Sup Fig 5). In addition, the co-occurrence of *tnpA*_*REP*_ genes in a genome with *procA* genes is nearly universal. Among the 618 genomes of *Prochlorococcus*, the TnpA_REP_ are only found within the LLIV clade (28 genomes), like prochlorosins [1]. Except for a few exceptions, likely due to genome incompleteness, if a genome encodes prochlorosins it has *tnpA*_*REP*_ genes, and vice versa (Fig 1B).

Moreover, the arrangement of *tnpA*_*REP*_ (full-length and truncated fragments) and *procA* genes shows a clear pattern of association: First, the genes are always facing each other in opposite directions (see brackets in Fig 1A, Fig 2), while *procA* genes are always in *cis* with each other, and often regularly spaced in clusters [1]. The distance between the facing *tnpA*_*REP*_ genes and the facing *procA* is extremely well conserved across distantly related strains (Fig 2B, Sup Fig 5). This conservation of intergenic distance is unlikely to happen by chance and suggests a mechanism physically linking the genes, possibly through mobilization by the transposase.

**FIG 2.**
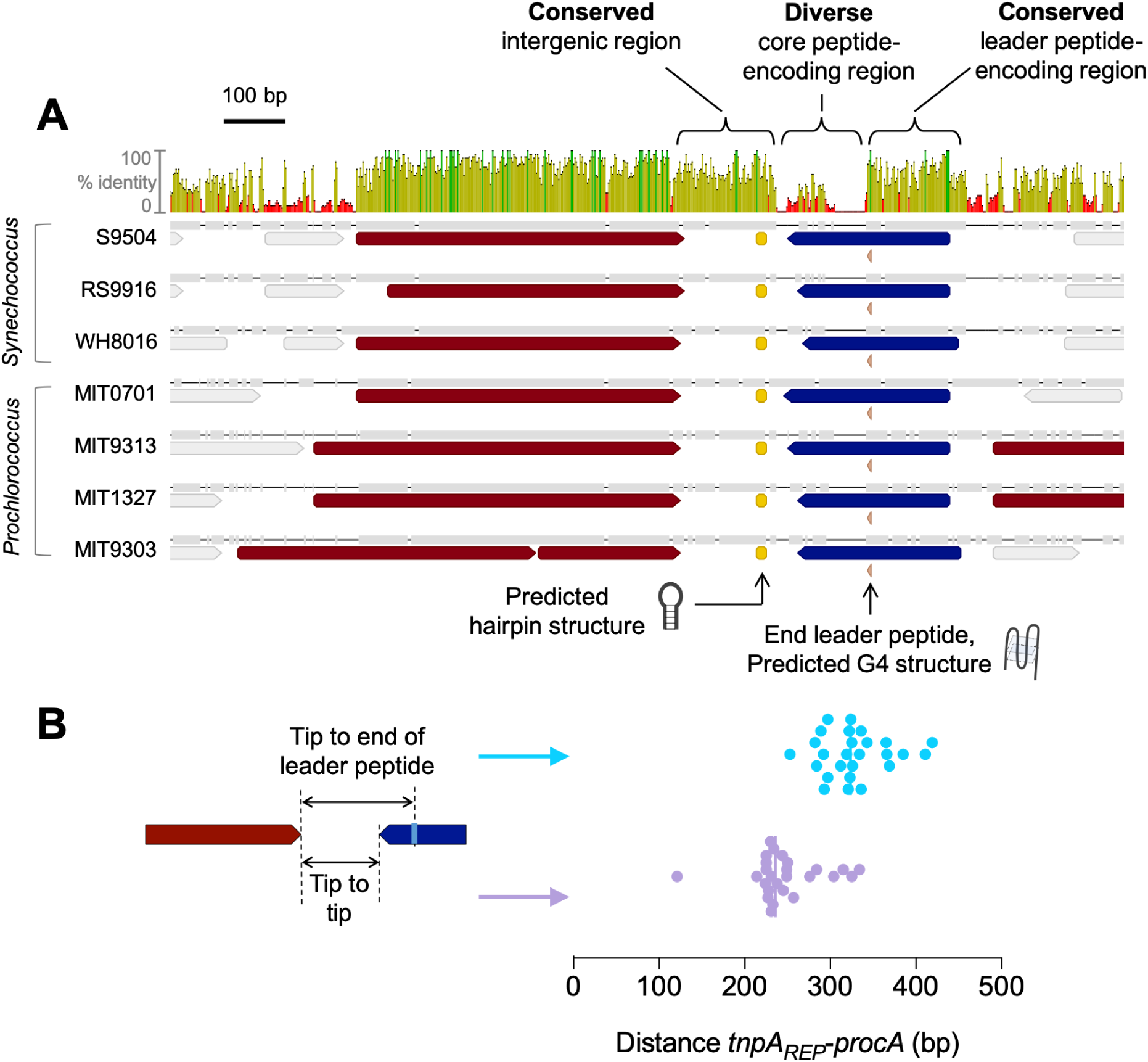
**(A)** Alignment of selected genomic regions containing full-length *tnpA*_*REP*_ (red) facing *procA* genes (blue) across the phylogeny of marine picocyanobacteria. The upper panel shows the percentage of nucleotide sequence identity (colors for each given position are green for 100% identity, yellow between 30 - 100%, red below 30%). The border between leader to core peptide region is indicated (triangle at the end of the leader peptide-encoding region). As previously described [1] the leader peptide-encoded region of *procA* is conserved across strains while the core peptide-encoding region is highly variable and contains large numbers of indels. Interestingly, in the case of facing *tnpA*_*REP*_ and *procA* genes, the *tnpA*_*REP*_ and the intergenic region are equally well conserved, showing that diversity is precisely focused on the core peptide-encoding region as the gene pair evolves. Two types of DNA secondary structures are predicted to form (see methods for detail): a ‘hairpin’ structure in the intergenic region, and a guanine quadruplex (G4) structure exactly at the end of the leader peptide region. **(B)** Distance between *tnpA*_*REP*_ and *procA* gene pairs in all 26 instances in the data set (see Sup Fig 5) where the genes face each other. Since the core peptide-encoding region tends to vary in length, both distance from the tip of *tnpA*_*REP*_ to the tip of facing *procA* (purple) and to the end of the leader peptide of facing *procA* (turquoise) are shown.

Second, the intergenic sequence between the facing *tnpA*_*REP*_ and *procA* genes is well-conserved among marine *Synechococcus* to *Prochlorococcus* genomes, as well conserved as the leader peptide-encoding sequence (Fig 2A), suggesting that it plays a role in the association.

To begin to unravel the mechanism of association we searched for potential TnpA_REP_ recognition targets. Of note, marine picocyanobacteria belong to a TnpA_REP_ subgroup that was not found to associate with REP-like sequences, but rather with inverted repeats (consensus sequence GGGG[AT][CG]A[CG]) [24], [25]. We could not find any sign of association with inverted repeats, but we did detect motifs forming DNA secondary structures: a hairpin structure (Fig 2A) in the intergenic sequence, and a Guanine quadruplex (G4) structure (Fig 2A) (see methods for details); both of these structure types have been shown to be potential recognition targets for TnpA_REP_ transposases [16], [17], [23]. Without further clues, the hairpin structure could simply correspond to conserved rho-independent terminators, which are common in cyanobacteria.

The G4 structures are more conspicuous as they form around a highly conserved GGCGG motif which delineates the transition from the leader to the core peptide. This site serves as an anchor point for the ProcM modifying enzyme [2]. Despite being a feature of only some of the *procA* genes (G4 is predicted to form in 25 to 45% of *procA* genes according to different prediction software, see methods for details), the conservation of occurrence at this particular site suggests that it could play a role. Interestingly, G4 structures have been shown to participate in pilin antigenic variation - one of the best described diversifying recombination mechanisms - in different proteobacteria such as *N. gonorrhoeae* [26]. However, the configuration in the latter is significantly different, as there the G4 is placed a few hundred base pairs upstream of the recombination hotspot, and recombination occurs by swapping a domain of the active pilin locus with the domain of idle pilin loci in the genome. In contrast, prochlorosins don’t result from domain swapping, but rather from indels in the core peptide region [1].

Overall, these results show that *tnpA*_*REP*_ and *procA* genes are physically linked in picocyanobacterial genomes in a non-random pattern, and this linkage is most probably the result of the TnpA_REP_ enzyme activity. Importantly, the closest TnpA_REP_ relatives in cyanobacteria (Sup Fig 1) do not colocalize with lanthipeptide-related genes, suggesting that this association is unique to marine picocyanobacteria resulting in the unique multiplication and diversification of lanthipeptide genes. Thus, we propose that similar to how TnpA_REP_ drives the expansion and diversification of the REP sequences repertoire in other bacterial genomes, the TnpA_REP_ homologs in marine picocyanobacteria genomes have associated with lanthipeptide genes, contributing to their expansion and diversification and resulting in the extreme diversity we see among prochlorosins. We suggest that the *tnpA*_*REP*_ / *procA* association enables a powerful diversifying recombination mechanism, precisely focused on modifying the core peptide encoding sequence, while maintaining the leader peptide sequence unchanged. We note that another model for the driving force of prochlorosin diversity has been proposed by Cubillos-Ruiz (2015) [27]. In that model, a putative *chi* site within the precursor peptide is postulated to serve as a hotspot for localized homologous recombination, mediated by the RecBCD DNA repair pathway. The two models are not necessarily mutually exclusive and could work in concert.

Testing the *chi* site hypothesis would involve demonstrating the chi site activity, whereas validation of the model we propose would require experimental assays determining the recognition sequence and catalytic activity of a *Prochlorococcus* TnpA_REP_ transposase as performed for the TnpA_REP_ homologs in *E. coli* [16], [17].

## MATERIALS AND METHODS

### Dataset

The genomes of *Prochlorococcus* LLIV clade and marine *Synechococcus* were taken from the IMG/ProPortal database (https://img.jgi.doe.gov/cgi-bin/proportal/main.cgi) hosted within the JGI Integrated Microbial Genomes system. A few additional *Synechococcus* genomes from National Center for Biotechnology Information (NCBI) database were added to the dataset: WH8101, UW179A, UW105, BS56D, and the metagenome-assembled genome EAC657. The set of *Prochlorococcus* and *Synechococcus* genomes containing *tnpA*_*REP*_ and/or *procA* genes that were used in this study are listed in Table S1.

### Gene annotation, sequence alignments, and phylogenetic tree construction

Prochlorosin annotations were taken from [1], and TnpA_REP_ from [18], [24]. Prediction of *procA* and *tnpA*_*REP*_ genes was expanded to new strains using BLASTP (E value of 1e-5) with ProcA or TnpA_REP_ protein sequence query. For Prochlorosins, the end of the leader peptide was predicted by searching the motif GCTGG[TAGC]GG in the *procA* sequence. For strain phylogeny, we used the built-in IMG-Proportal phylogeny tool and annotated the tree (Fig 1B) using the Interactive Tree Of Life (iTOL). The facing *procA-tnpA*_*REP*_ nucleotide sequence alignment (Fig 2A) was performed in Geneious Prime 2020.0.5 using the geneious aligner. The TnpA_REP_ protein sequence alignment was performed using MUSCLE and the tree (Sup Fig 1) was built using the Geneious tree builder (Neighbour-Joining); Only the TnpA_REP_ sequences above 200 amino acids were kept for the alignment. The *Prochlorococcus* MIT9313 TnpA_REP_ fragments protein sequence alignment (Sup Fig 4) was performed using the Geneious aligner.

### DNA secondary structure prediction

Hairpin structure formation was predicted using RNAstructure [28]. The consensus sequence ACTCA**AAGCCCTTGCATTAGCAGGGGCTT**TTTATT in the Fig 2A alignment intergenic region is predicted to form a hairpin structure (bold nucleotides) with a free energy of -10.3 kcal/mol.

Guanine quadruplexes (G4) were assessed using the dataset of 2509 distinct *procA* genes (Operating Prochlorosin Units, OPU) from [1]. In total, the G4-PREDICTOR V.2 software predicts G4 structures in 45% of OPUs, while the G4hunter software predicts G4 structures in 25.7% of OPUs.

### 3D structure modeling

The *in silico* prediction of *Prochlorococcus* MIT9313 PMT_0830 TnpA_REP_ 3D structure was performed using SWISS-MODEL, with the *E. coli* TnpA_REP_ (PDB_ID 4ER8) serving as the template for homology modeling.

## Supporting information

SFig 3

## ACKNOWLEDGMENTS

This study was supported in part by the Simons Foundation (Life Sciences Project Award IDs 337262, 509034SCFY17, 647135 and SCOPE Award ID 329108, to SWC), and an NSF-EDGE grant (1645061 to SWC).

We gratefully thank Yves Quentin for his detailed review of the manuscript and helpful suggestions.

