## Supplementary data for "Association of lanthipeptide genes with TnpA_REP_ transposases in marine picocyanobacteria"

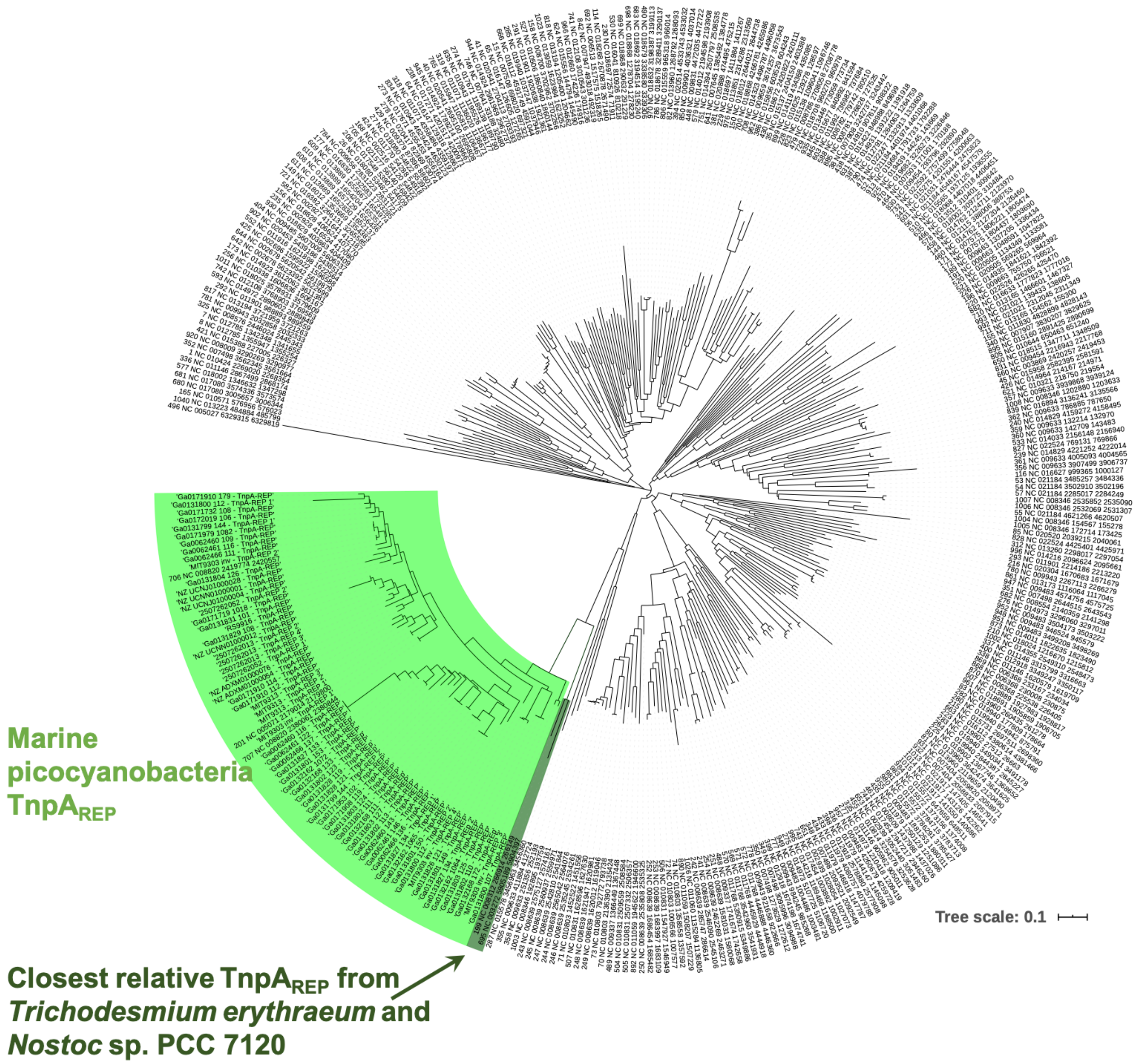


**Sup Fig 1 -** Phylogenetic tree of the marine picocyanobacteria $\text{TnpA}_{\text{REP}}$ homologs identified in this study, within the $\text{TnpA}_{\text{REP}}$ group in which they cluster from [1] (RAYT group 1). The closest relative homolog $\text{TnpA}_{\text{REP}}$ are found in the genomes of cyanobacteria *Trichodesmium erythraeum* (strain IMS101) and *Nostoc* sp. PCC 7120 which are not co-localized with lanthipeptide genes on the genome.

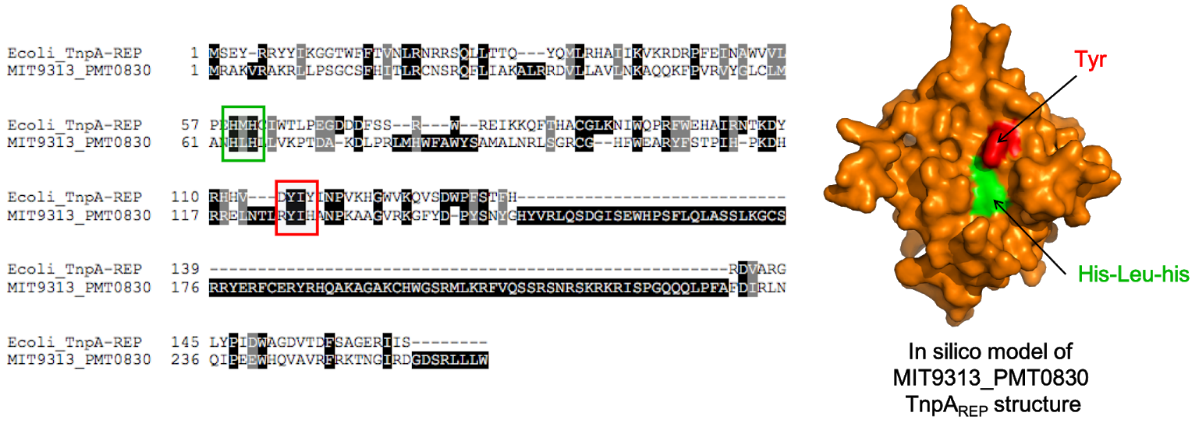


**Sup Fig 2 - (A)** Alignment of MIT9313 TnpA$_{REP}$ with the homolog transposase from *E. coli [2]* for which a 2.6 Å crystal structure is available, generated using T-coffee [3]. Active site residues are highlighted by the green and red boxes. The *Prochlorococcus* TnpA$_{REP}$ possesses an additional domain on its C-terminal side compared to the *E. coli* homolog. **(B)** 3D model of *Prochlorococcus* MIT9313 TnpA$_{REP}$ (PMT_0830) generated using Swiss-Model [4], highlighting the predicted transposase active site residues.

**Sup Fig 3** - All $\mathit{tnpA}_{REP}$ occurrences in marine picocyanobacteria used in this study. Red gene annotations correspond to predicted $TnpA_{REP}$ homologs, blue annotations are predicted *procA* genes. All other genes are in grey. Note that most occurrences of $\mathit{tnpA}_{REP}$ are from genomes sequenced from a single-cell [5] which are incomplete and fragmented. Contigs often terminated next to transposases genes, probably due to the difficulty of assembling these repeat-containing regions. Thus the co-localization of $\mathit{tnpA}_{REP}$ with *procA* genes across the dataset is an underestimation.

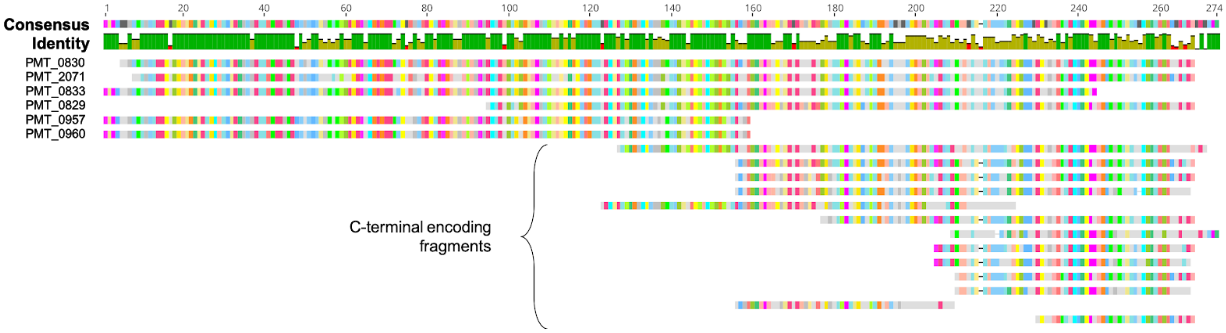


**Sup Fig 4** - Alignment of all $\mathit{tnpA}_{REP}$ occurrences in *Prochlorococcus* MIT9313 genome. Multiple copies are most probably non-functional degraded fragments of the transposase, often corresponding to the C-terminal extremity of the protein. While most fragments correspond to truncated ORFs, a Tblastn search reveals that a few fragments result from frameshifts in the coding sequence.

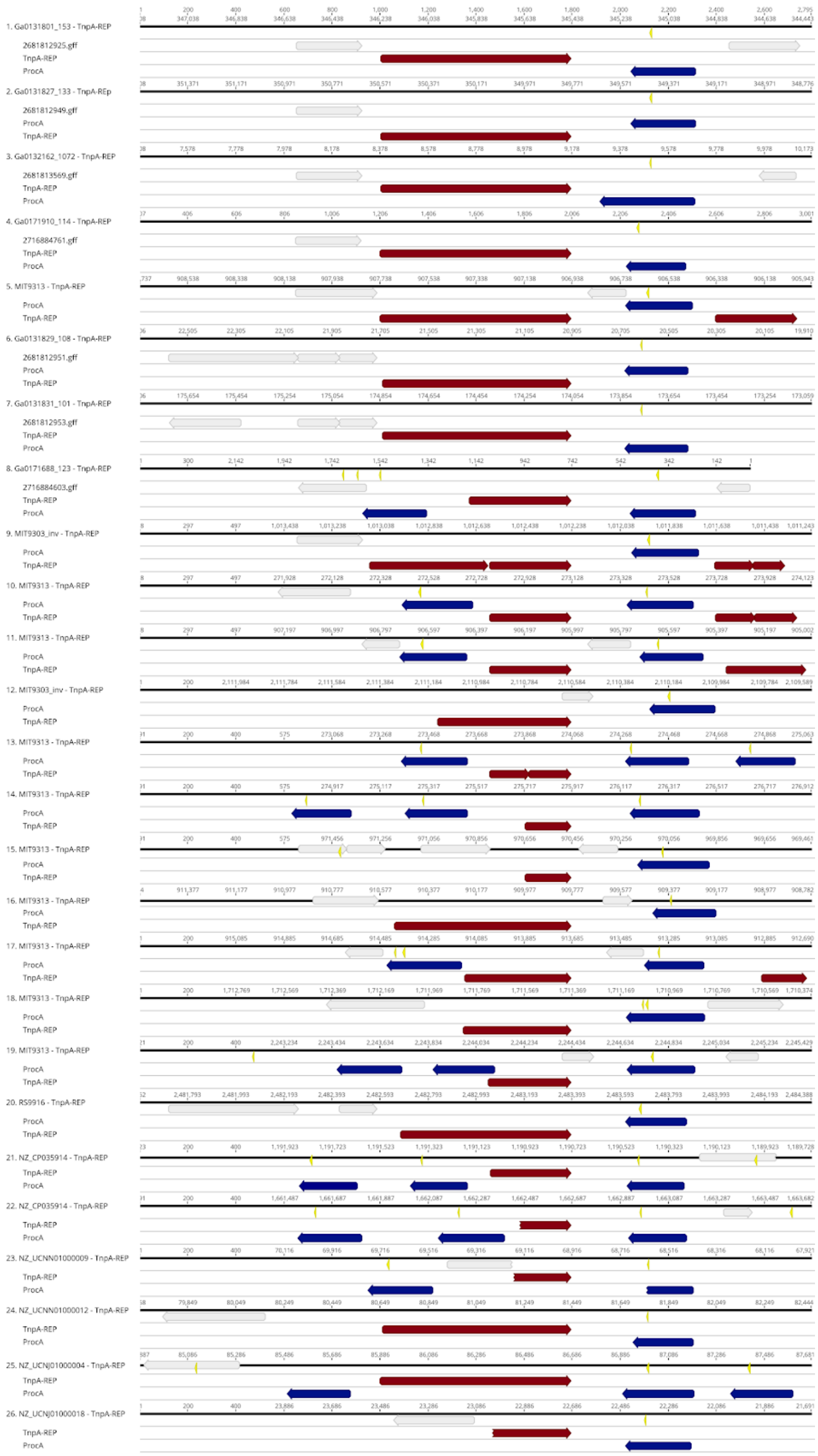


**Sup Fig 5** - All *tnpA*$_{REP}$ / *procA* facing pairs used for Fig 2B. Red gene annotations correspond to predicted TnpA$_{REP}$ homologs, blue annotations are predicted *procA* genes (see methods for details). Short yellow annotations show the predicted end of *procA* leader peptide-encoding region. All sequences have been aligned on the end of the *tnpA*$_{REP}$ gene.

**Table S1** - *Prochlorococcus* and *Synechococcus* genomes used in this work, reporting the number of $TnpA_{REP}$ and ProcA Blastp hits (aa: amino acids).

| Genome ID | Genome name | Number of $TnpA_{REP}$ hits (>200 aa) | Number of $TnpA_{REP}$ fragment hits (<200 aa) | Number of ProcA hits |
|---|---|---|---|---|
| 637000211 | *Prochlorococcus marinus* MIT9313 | 3 | 10 | 32 |
| 639857007 | *Synechococcus* sp. RS9916 | 1 | 0 | 22 |
| 640069323 | *Prochlorococcus marinus* MIT9303 | 3 | 14 | 26 |
| 640427149 | *Synechococcus* sp, WH7803 | 0 | 0 | 2 |
| 649990023 | *Synechococcus* sp. CB0205 | 2 | 1 | 0 |
| 649990022 | *Synechococcus* sp, CB0101 | 0 | 0 | 7 |
| 2507262013 | *Synechococcus* sp. KORDI-100 | 4 | 0 | 11 |
| 2507262052 | *Synechococcus* sp. WH 8016 | 2 | 1 | 3 |
| 2606217681 | *Prochlorococcus* sp. MIT0702 | 3 | 20 | 19 |
| 2606217682 | *Prochlorococcus* sp. MIT0703 | 3 | 14 | 19 |
| 2606217684 | *Prochlorococcus* sp. MIT0701 | 3 | 17 | 19 |
| 2681812923 | *Prochlorococcus* sp. MIT1303 | 2 | 4 | 5 |
| 2681812924 | *Prochlorococcus* sp. MIT1306 | 2 | 2 | 7 |
| 2681812925 | *Prochlorococcus* sp. MIT1312 | 3 | 5 | 17 |
| 2681812926 | *Prochlorococcus* sp. MIT1313 | 4 | 2 | 5 |
| 2681812927 | *Prochlorococcus* sp. MIT1318 | 3 | 3 | 5 |
| 2681812928 | *Prochlorococcus* sp. MIT1320 | 1 | 2 | 7 |
| 2681812948 | *Prochlorococcus* sp. MIT1323 | 1 | 6 | 4 |
| 2681812949 | *Prochlorococcus* sp. MIT1327 | 3 | 4 | 17 |
| 2681812950 | *Prochlorococcus* sp. MIT1342 | 2 | 4 | 8 |
| 2681812951 | *Synechococcus* sp. MIT9504 | 1 | 0 | 88 |
| 2681812952 | *Synechococcus* sp, MIT9508 | 0 | 0 | 9 |
| 2681812953 | *Synechococcus* sp. MIT9509 | 1 | 0 | 89 |
| 2681813566 | *Prochlorococcus* sp. MIT1205 | 0 | 6 | 4 |
| 2681813569 | *Prochlorococcus* sp. MIT1227 | 3 | 4 | 17 |
| 2681813575 | *Prochlorococcus* sp. MIT1418 | 3 | 5 | 5 |
| 2716884257 | Uncultured *Synechococcus* sp, AG-673-D02 | 0 | 0 | 8 |
| 2716884268 | Uncultured *Prochlorococcus* sp. AG-436-F13 | 0 | 3 | 7 |
| 2716884270 | Uncultured *Prochlorococcus* sp, AG-436-J08 | 0 | 0 | 2 |
| 2716884427 | Uncultured *Prochlorococcus* sp. AG-432-D09 | 1 | 0 | 4 |
| 2716884444 | Uncultured *Prochlorococcus* sp. AG-436-P23 | 1 | 3 | 9 |
| 2716884464 | Uncultured *Prochlorococcus* sp. AG-463-F02 | 1 | 3 | 3 |
| 2716884603 | Uncultured *Synechococcus* sp. AG-323-O22 | 0 | 2 | 15 |
| 2716884611 | Uncultured *Synechococcus* sp, AG-670-D07 | 0 | 0 | 1 |
| 2716884612 | Uncultured *Synechococcus* sp. AG-670-F04 | 0 | 2 | 3 |
| 2716884625 | Uncultured *Synechococcus* sp. AG-683-A02 | 1 | 1 | 4 |
| 2716884630 | Uncultured *Synechococcus* sp. AG-686-A03 | 0 | 1 | 0 |
| 2716884679 | Uncultured *Synechococcus* sp. AG-686-D09 | 1 | 0 | 4 |
| 2716884758 | Uncultured *Prochlorococcus* sp. AG-412-I05 | 0 | 9 | 8 |
| 2716884759 | Uncultured *Prochlorococcus* sp. AG-412-I20 | 1 | 2 | 3 |
| 2716884761 | Uncultured *Prochlorococcus* sp. AG-412-L10 | 3 | 0 | 9 |
| 2716884620 | Uncultured *Synechococcus* sp, AG-676-C06 | 0 | 0 | 4 |
| 2716884666 | Uncultured *Synechococcus* sp, AG-673-B04 | 0 | 0 | 4 |
| 2716884669 | Uncultured *Synechococcus* sp, AG-679-A04 | 0 | 0 | 27 |
| 2716884671 | Uncultured *Synechococcus* sp, AG-679-C18 | 0 | 0 | 7 |
| BS56D | *Synechococcus* sp. BS56D | 1 | 0 | 3 |
| EAC567 | Uncultured *Synechococcus* sp. EAC567 | 0 | 0 | 16 |
| NZ_CP035914 | *Synechococcus* sp. WH 8101 | 0 | 2 | 18 |
| UW105 | *Synechococcus* sp. UW105 | 1 | 2 | 12 |
| UW179A | *Synechococcus* sp. UW179A | 1 | 3 | 51 |
