## Supplementary figures and images for "Association of lanthipeptide genes with TnpA_REP_ transposases in marine picocyanobacteria"

### SFig 3

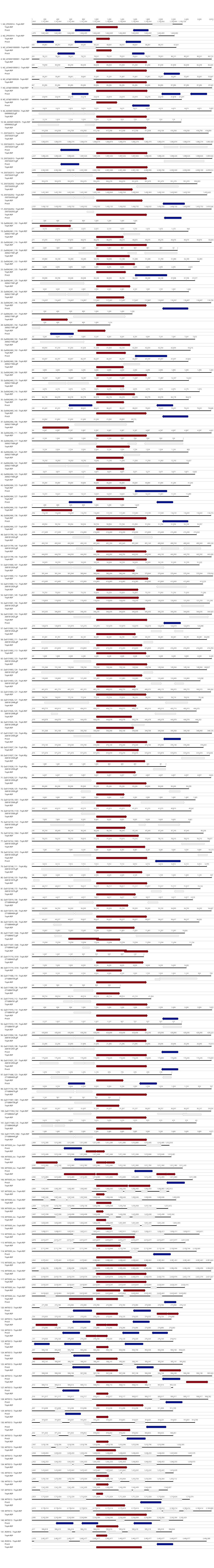
